# Fetal maturation revealed by amniotic fluid cell-free transcriptome in rhesus macaques

**DOI:** 10.1101/2021.04.28.441815

**Authors:** Augusto F. Schmidt, Daniel Schnell, Kenneth Eaton, Kashish Chetal, Paranthaman S. Kannan, Lisa A. Miller, Claire A. Chougnet, Daniel T. Swarr, Alan H. Jobe, Nathan Salomonis, Beena D. Kamath-Rayne

## Abstract

Accurate estimate of fetal maturity could provide individualized guidance for delivery of complicated pregnancies. However, current methods are invasive, have low accuracy, and are limited to fetal lung maturation. To identify diagnostic gestational biomarkers, we performed transcriptomic profiling of lung and brain, as well as cell-free RNA from amniotic fluid of preterm and term rhesus macaque fetuses. These data predict new and prior associated gestational age differences in distinct lung and neuronal cell populations when compared to existing single-cell and bulk RNA-Seq data. Comparative analyses found over 200 genes coincidently induced in lung and amniotic fluid, and dozens in brain and amniotic fluid. This data enabled creation of computational models that accurately predicted lung compliance from amniotic fluid and lung transcriptome of preterm fetuses treated with antenatal corticosteroids. Cell-free RNA in amniotic fluid may provide a substrate of global fetal maturation markers for personalized management of at-risk pregnancies.

## Introduction

Preterm birth is responsible for 1 million infant deaths every year worldwide, with an even larger number of infants surviving with long-term respiratory and neurodevelopmental morbidities (Blencowe et al., 2013). Accurate assessment of fetal lung maturation could help in guiding timely delivery of complicated pregnancies and administration of antenatal therapies that prevent morbidities associated with preterm birth, such as antenatal corticosteroids (ACOG, 2021; Spong et al., 2011). However, the traditional methods of assessment of fetal maturation are invasive, limited to the evaluation of fetal lung maturation and do not provide information on maturational state of other fetal organs (Geary and Whitsett, 2005; Kamath-Rayne and DeFranco, 2014). Maturational state of the fetal brain is particularly important given the continued growth and development of this organ through the third trimester and the increased risk of neurodevelopmental disabilities in preterm infants, even among those delivered in the late preterm period (34 to 36 weeks’ gestation) (Kugelman and Colin, 2013). Fetal maturity can be also be accelerated clinically by administration of antenatal corticosteroids, which has been standard of care for pregnancies at risk of preterm delivery for more than 30 years (ACOG, 2021). Antenatal corticosteroids have pleiotropic positive effects in the preterm fetus, particularly to the lung, but also potentially detrimental effects, particularly to brain development (Moisiadis and Matthews, 2014; Schmidt et al., 2020; Schmidt et al., 2019a). Fetal maturity testing that provides information on global maturity and the ability to longitudinally track fetal development and response to treatment would be an ideal tool for management of high-risk pregnancies and delivery planning.

To overcome the challenge of determining global fetal maturation, we evaluated the cell-free transcriptome of amniotic fluid for existing and novel markers of fetal lung and brain maturation in a nonhuman primate model. Amniotic fluid contains fetal cell-free DNA, RNA, and proteins, and is a reflection of fetal status (Underwood et al., 2005; Zwemer and Bianchi, 2015). Furthermore, the composition of the amniotic fluid changes according to gestational age and fetal conditions (Hui et al., 2012; Hui et al., 2013; Jang et al., 2017; Kamath-Rayne et al., 2014; Vora et al., 2017). Analyses of human pregnancies show that a portion of the fetal cell-free RNA in amniotic fluid can be traced to genes and biological processes associated with systems and organs that are not in direct contact with the amniotic fluid, such as the central nervous system (Kamath-Rayne et al., 2015). Moreover, the transcriptome of amniotic fluid changes with advancing gestation, suggesting the presence of markers of fetal maturity (Kamath-Rayne et al., 2015). Cell-free RNA is of particular interest due to its presence not only in the amniotic fluid but also, in smaller amounts, in the maternal serum. Circulating RNA correlates with changes in maternal-fetal development, as well as neurological diseases (Koh et al., 2014; Ngo et al., 2018). Analyses of the cell-free RNA in amniotic fluid could provide a foundation for targeted testing of fetal maturation markers in the maternal serum. While data from human samples raise intriguing questions regarding the presence of tissue-specific transcripts in amniotic fluid, these studies cannot directly evaluate transcriptome changes in fetal organs.

In this context, the amniotic fluid could provide information on the effects of antenatal corticosteroids beyond the lung and guide personalized treatment based on individual assessment of the maturational stage of various fetal organ systems. In a rhesus macaque model, we tested the hypothesis that changes to the amniotic fluid transcriptome reflect gene expression changes in the fetal lung and the fetal brain, which correlate with intended effects (promotion of fetal maturity) or unintended effects from antenatal corticosteroid treatment.

## Results

### Fetal maturation is associated with broad coordinated transcriptional programs in lung, brain and amniotic fluid

We used a rhesus macaque model of fetal maturation to identify genes that are developmentally regulated from preterm to term gestation and by the effect of antenatal corticosteroids in the amniotic fluid, fetal lung, and fetal brain. Time-mated and ultrasound dated pregnant rhesus macaque fetuses were delivered with intact membranes by cesarean section at preterm (∼132 days) or near-term (∼155 days) gestation, with full term being 165 days of gestation. The gestational age of delivery of preterm animals corresponds to about 32 weeks’ gestation in human, when the lungs are at the saccular stage of development. Other groups of animals were treated with different formulations and doses of antenatal corticosteroids 5 days prior to delivery at preterm gestation (∼132 days) (**Fig. 1A, Supplemental Table 1**). For each control time-point and antenatal corticosteroid treatment, multiple replicates were collected for matched fetal lung, fetal brain (hippocampus) and amniotic fluid; these were profiled using RNA-Seq, to allow for both differential and sample-level correlative analyses. As the specimens were collected over multiple mating seasons, batch effects correction methods were applied to the resulting data, considering fetal sex, season, and treatment group, following gene-level transcript per million (TPM) quantification. When visualized using Principal Component Analysis, the transcriptomes of nearly all samples were readily separable by treatment groups and time-points, with most intermixing in hippocampus, as expected (**Fig 1B**). As these data suggest that each treatment regimen results in distinct molecular impacts, we further examined the presence of unique marker genes and enriched gene-sets for each treatment group in lung, hippocampus and amniotic fluid. Indeed, we found clear gene modules that defined each treatment group and term gestation in the fetal lung, with maturity at term most indicated by development of innate immunity (neutrophil activation, NADPH oxidase) (**Fig. 1C, Supplemental Table 2**). Comparison of this maturity gene expression signature showed expected heterogeneity among the treatment groups, with less variability in preterm controls (**Fig. 1C**, bottom cluster). Likewise, in hippocampus, we found term maturation to be associated with myelination of neurons, with antenatal corticosteroid treatment differentially impacting Golgi-processing pathways (**Supplemental Fig. 1, Supplemental Table 2**). Global comparison of gene expression in term and corticosteroid-treated samples relative to preterm controls indicated dominant expression differences at term in lung and brain, while in amniotic fluid most differences were seen in PO Dexamethasone treated macaques (**Fig. 1D, Supplemental Table 3**). Interestingly, while amniotic fluid showed a trend towards predominant upregulation in gene expression at term, brain showed the opposite pattern.

**Figure 1.**
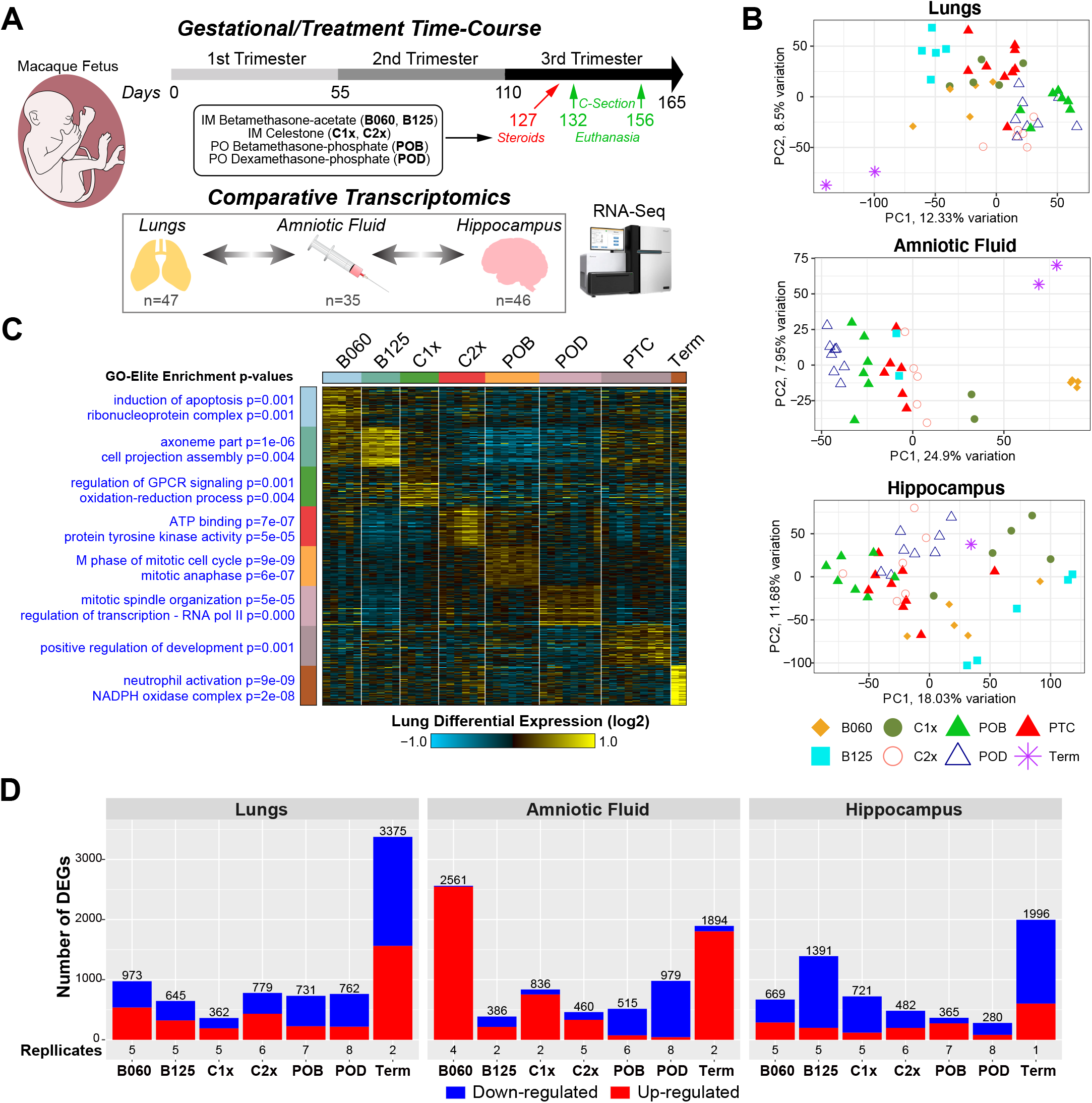
Evaluation of antenatal corticosteroids across tissues and amniotic fluid during rhesus gestation. A) Study design for treatment and c-section of pregnant Rhesus Macaque females using indicated antenatal corticosteroid dosing. Amniotic fluid, brain (hippocampus) and lung tissue were collected from each fetus from either pre-term (127-136 days) or term (156 and 157) gestation and analyzed by bulk RNA-Seq analysis. B) Principal Component Analysis of all genes, following NOISeq batch-effect correction and, for Amniotic fluid, brain and lungs (mean TPM≥1). C) Heatmap of the top most specific marker genes for lung RNA-Seq for each treatment. Expression values were calculated as log2 fold-changes relative to the mean of each row. The top GO-Elite Gene Ontology enrichment results are denoted to the left of each cluster along with corresponding Fischer’s Exact test enrichment p-values. D) Number of differentially expressed genes in all treatments to pre-term controls (fold ≥ 1.25 and limma p-value ≤ 0.05 adjusted for treatment year effects). B060: intramuscular (IM) betamethasone-acetate 0.06mg/kg x1 dose, B125: IM betamethasone-acetate 0.125mg/kg x1 dose, C1x: IM Celestone® (betamethasone-acetate+betamethasone-phosphate) 0.25mg/kg x1 dose, C1x: IM Celestone® 0.25mg/kg x2 doses, POB: oral betamethasone-phosphate 0.15mg/kg x3 doses, POD: oral dexamethasone-phosphate 0.15mg/kg x3 doses, PTC: preterm control.

### Gestation gene-expression differences are associated with cell-type and tissue-specific cell-free RNAs

While the rhesus macaque samples were profiled using bulk RNA-Seq, we investigated whether we could assess the contribution of specific cell-types and associated mRNAs for each of the tissue/amniotic datasets. To this end, we identified markers for previously defined discrete cell-populations from two large human single-cell (sc) RNA-Seq compendiums (lung and brain) as well as a large baboon pan-tissue bulk RNA-Seq library, as a reference for cell-type frequency prediction (deconvolution) (see Methods). Our *in silico* analysis of the potential tissue-of-origin for cell-free RNAs in amniotic fluid suggested that the most frequent cellular sources were esophagus, hypothalamus, pancreas, lower-leg muscle, skin, and lungs (**Fig. 2A, Supplemental Table 4**). As placenta samples were not a part of this large reference baboon tissue compendium (Mure et al., 2018), it is possible that the hypothalamus, a major source of neurohormones, was a substitute for placenta hormone secreting cells in this analysis. In amniotic fluid, term samples showed similar RNA contribution from the different organs as the preterm samples, with the exception of esophagus, which had highly represented RNA at term, while pancreas showed decreased associated RNAs at term. For rhesus macaque lung compared to a recent human lung scRNA-Seq compendium neonates through adults (Wang et al., 2020), we observed the largest contribution of mRNAs from matrix fibroblasts, erythrocytes, alveolar type 1 (AT1), ciliated cells, Capillary endothelial and natural killer (NK) cells. We noted that at term, there was a predominant increase in the predicted frequency of AT1 and alveolar type 2 (AT2) cell RNAs and a decrease in ciliated cells and vascular or airway smooth muscle cells (**Fig. 2B**). Cell-type predictions in the hippocampus, using a human pediatric cortex scRNA-Seq as a reference (Velmeshev et al., 2019), found that neuron, astrocyte, endothelial cell, and microglia represented the principal sources of hippocampal RNA. Term hippocampus RNAs were enriched in oligodendrocyte profiles, relative to all other gestational samples (**Fig. 2C**).

**Figure 2.**
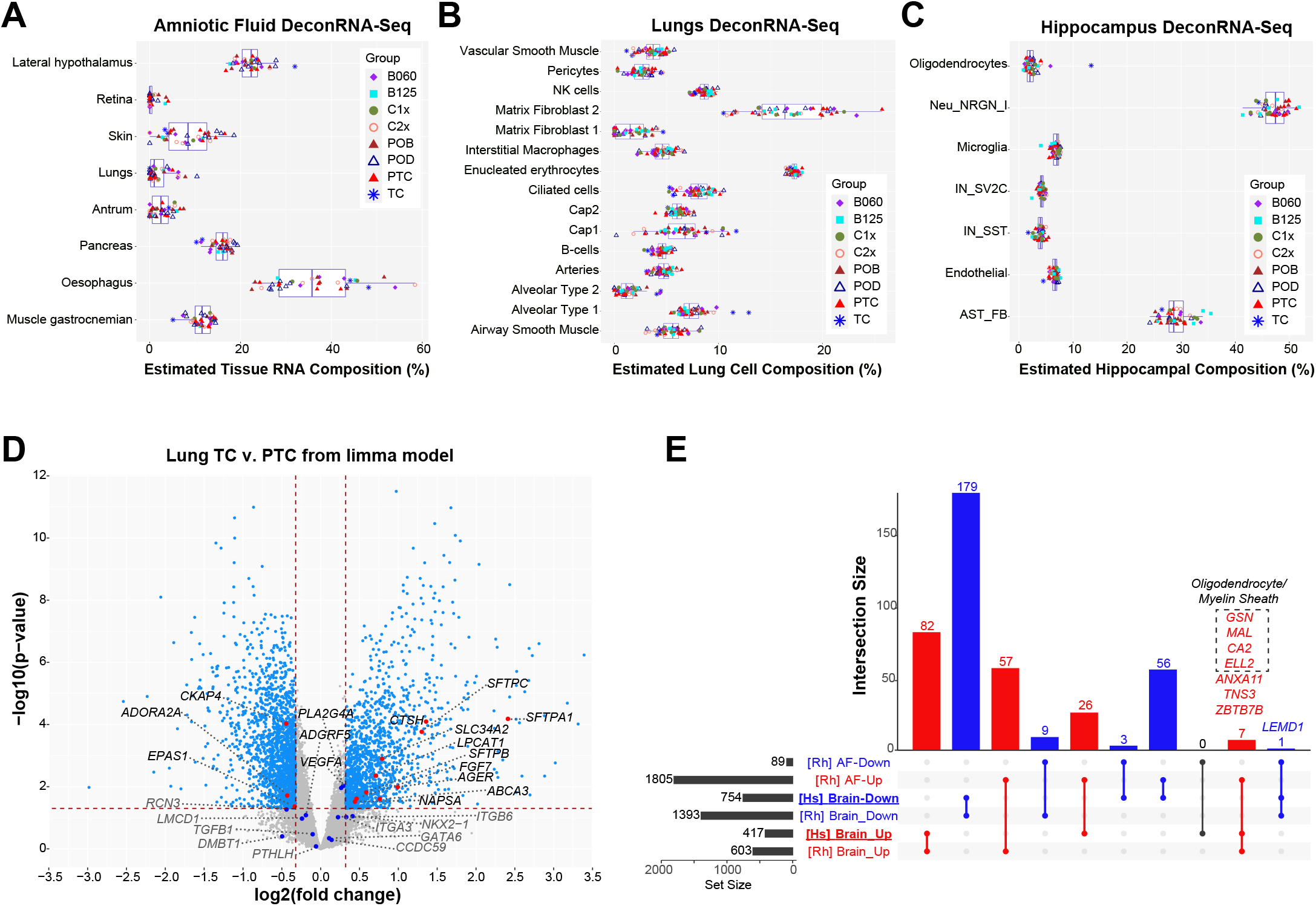
Established molecular markers and cell-types delineate term and pre-term samples. A) Predicted relative frequency of over 60 considered primate tissue regions in all evaluated Rhesus tissues from this study using the software DeconRNASeq. Only the most frequently detected tissues shown with specific estimates for all Rhesus samples by treatment group displayed. B) Predicted frequency of human lung cell-types, including neonatal, child and adult cell-types for only Rhesus lung RNA-Seq samples. C) Predicted frequency of brain cortex cell-types, collected from human pediatric and young adult, in all Rhesus brain RNA-Seq samples; deconvolution relative to single-cell RNA-Seq. D) Volcano plot of differentially genes comparing the limma p-value and fold-change for all genes in term amniotic fluid versus preterm controls. Prior defined human term induced genes from amniotic fluid, are highlighted red (significant) or blue (non-significant) (fold ≥ 1.25 and limma p-value ≤ 0.05 adjusted for treatment year effects). E) Upset plot indicating the overlap of differentially expressed genes in human neonatal versus mid-gestation hippocampus, compared to rhesus macaque term versus preterm hippocampus and/or amniotic fluid. Genes associated with Oligodendrocyte cell identify and myelin sheath formation are highlighted.

To determine whether transcripts identified as uniquely induced in term corresponded to previously identified markers of fetal maturation, we compared all differentially expressed genes in lung and amniotic fluid to our prior identified set of human term amniotic fluid induced transcripts (Kamath-Rayne et al., 2015). In the rhesus macaque lung, 10 of 25 human amniotic fluid markers were overexpressed at term compared to preterm, several of which are well-defined markers of lung maturation (*SFTPA1, SFTPC, SFTPB, LPCAT1, FGF7, CTSH)* (**Fig. 2D**). Although only one of these transcripts met the established significance threshold in macaque amniotic fluid, with the remainder just below the statistical threshold for significance, nearly all such markers matched the same direction of regulation (**Supplemental Fig. 2A**). To validate term induced macaque hippocampal transcripts, we compared these data to differentially expressed genes in human neonate/infant versus preterm (21 and 24 weeks) hippocampus, using bulk RNA-Seq from the Allen Brain Atlas (**Supplemental Table 5**). While only one viable term hippocampus sample was analyzed from the rhesus macaque, we observed highly specific gene expression overlaps from human and rhesus macaque, with 82 upregulated and 179 downregulated genes conserved (**Fig. 2E**).

### Term gestation is associated with inflammatory signaling in the lung and amniotic fluid

We next looked for genes with shared patterns of differential expression between amniotic fluid and lung or brain that might serve as peripheral indicators of fetal organ maturation. In the brain, we identified 57 upregulated and 9 downregulated genes with same pattern of expression in term versus preterm. Among these, 7 genes were upregulated in macaque and human brain as well as amniotic fluid (**Fig. 2E**). Intriguingly, the majority were associated with oligodendrocyte identity and myelin sheath formation (*GSN, MAL, CA2, ELL2*). In lungs we identified 228 genes commonly upregulated in amniotic fluid, and 12 genes commonly downregulated (**Supplemental Fig. 2B**). Gene-set enrichment of upregulated genes shared by amniotic fluid- and lung-term samples relative to the preterm controls showed an induction of mRNAs associated with inflammatory processes, in particular, monocyte activation, in addition to lung maturation (**Supplemental Fig. 3A**). Transcription factor binding site analysis predicted activation of IRF8, STAT3, SPI1 and NFKB/RELA, shared in amniotic fluid and lung, corroborating the activation of inflammatory pathways with term gestation in these tissues (**Supplemental Fig. 3B**). Overall, advancement from preterm to near term gestation showed activation of signaling of pro-inflammatory pathways and immune cell activation in the fetal lung and amniotic fluid.

As noted, the number of commonly deregulated genes in amniotic fluid is likely underestimated, given most prior established lung marker genes were just below the threshold of statistical significance. To account for this, we performed an orthogonal analysis, taking advantage of the fact that we have paired tissue and amniotic fluid from the same animals. Correlation of expressed genes in amniotic fluid and lungs or brain of all 104 analyzed samples yielded hundreds of correlated transcripts (**Fig. 3A, B** and **Supplemental Table 6)**. These included well-established lung maturation marker genes, including *SFTPC, SFTPA1, SFTA2, SCGB3A2* and others (**Fig. 3A, Supplemental Table 7**). Gene-set enrichment of lung-amniotic fluid correlated transcripts using a large collection of prior reported single-cell markers found NK cell, AT2, AT1, Clara, granulocyte, and proximal cells among the enriched cell types (**Fig. 3C**). Conversely, we found enrichment of markers of vasculature including muscle and endothelial cells among brain-amniotic fluid correlated transcripts (**Fig. 3D**).

**Figure 3.**
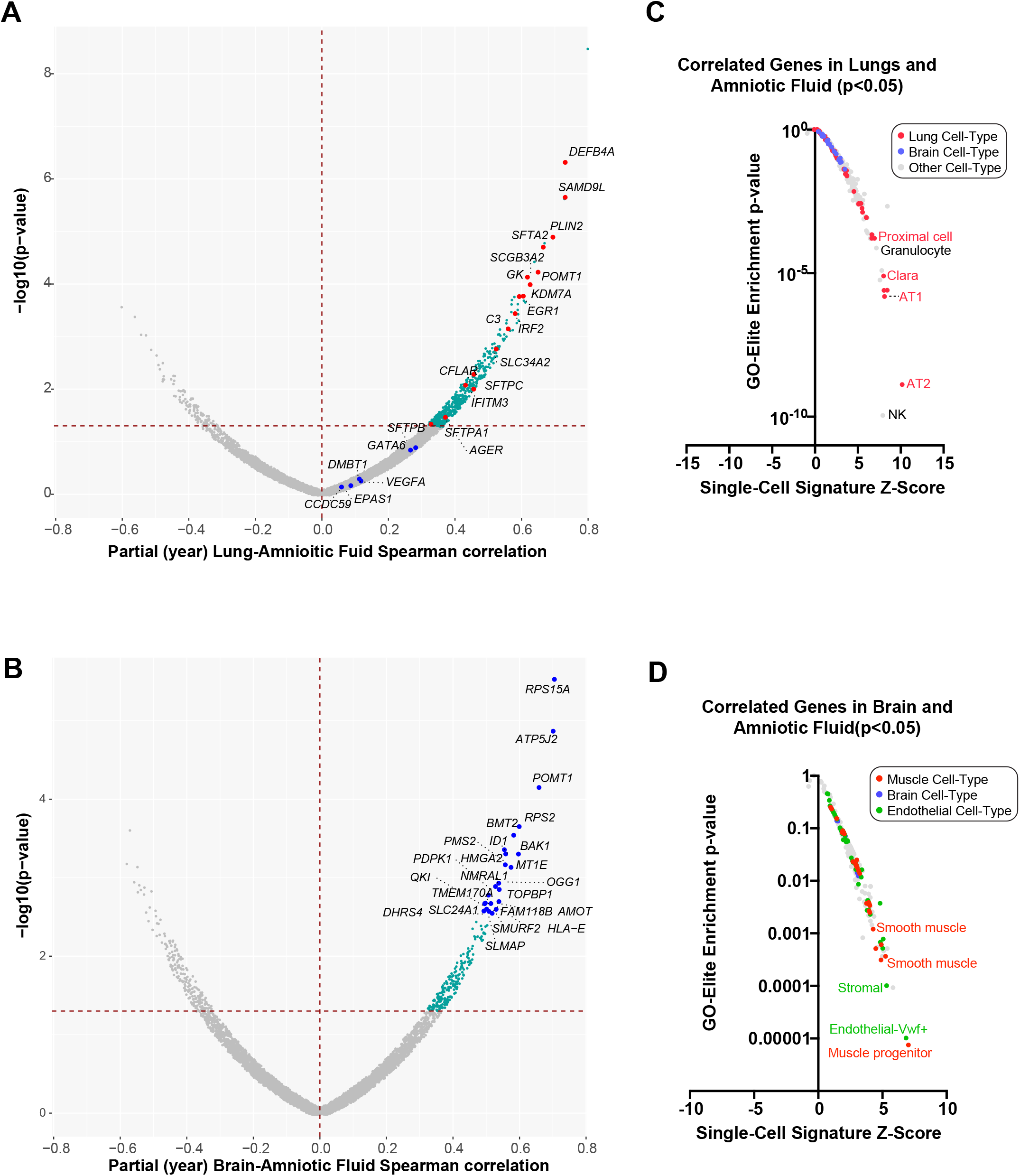
Detection of lungs and brain maturation programs in Amniotic Fluid. A-B) Scatter plot of highly correlated genes (Spearman Rank, controlling for treatment year) from the same animals (treatment and controls) in A) Amniotic Fluid and Lungs or B) Amniotic Fluid and Hippocampus. Spearman p-value and rank correlation are shown in the plot, with genes previously observed to be induced in human amniotic fluid, called out and designated by red (significant correlation) or blue (non-significant) in panel A. In panel B, the top most significant genes (p-value) are designated in blue. C-D) Gene-set enrichment of all available single-cell cell-type signatures for C) Amniotic-Fluid/Lung positively correlated transcripts (p≤0.05) or D) Amniotic-Fluid/Brain positively correlated transcripts. Gene-sets for Lung associated with Lung or Brain are specifically denoted in panel C and those for Muscle, Brain or Endothelial (most frequently enriched) are denoted in panel D.

### Amniotic fluid mRNAs can accurately predict fetal organ maturity

To evaluate maturity of fetal macaque lung, with and without antenatal corticosteroid treatment, we measured lung gas volume at 40 cmH_2_O on the pressure volume curve (PV40) **(Fig. 4A**). These data show that 25 out of the 36 treated animals had PV40 measurements greater than the range of preterm controls. Correlation of gene expression and PV40 found 727 genes in lungs, 234 genes in amniotic fluid, and 260 genes in brain that correlate with lung maturation (**Supplemental Table 8**). However, only a handful of organ-PV40 correlated genes overlapped between lung with amniotic fluid (*ATF1, CBLB, CEACAM1, GPD1L, KRAS, LARP4B, MAL2, PLIN2, SEC24A, SERTAD3*) or brain and amniotic fluid (*ADM, APPL2, ATF1, LYPLA1, NDUFA2, TMEM60*). Given that such gene programs likely reflect differing abundance of tissue resident and non-resident cell populations, we examined each of these broader gene sets using gene set enrichment against marker genes for diverse scRNA-Seq defined cell populations. Comparison of PV40 correlated transcripts in lung and amniotic fluid finds the common enrichment of mRNAs associated with monocyte, neutrophil, macrophage and NK-cells (**Fig. 4B, Supplemental Table 9**). When considering only lung enriched cell-type signatures, we observed a broad and specific enrichment of lung cell-type signatures, in particular of AT1 and AT2 cells. Similarly, comparison of PV40 correlated transcripts in brain and amniotic fluid finds the common enrichment of monocyte and NK-cell programs, in addition to B-cell, vascular endothelial and motor neutron programs (**Fig. 4C, Supplemental Table 9**). Brain-specific enriched PV40 correlated signatures were most frequently represented by neuron and glial cell markers along with endothelial cells.

**Figure 4.**
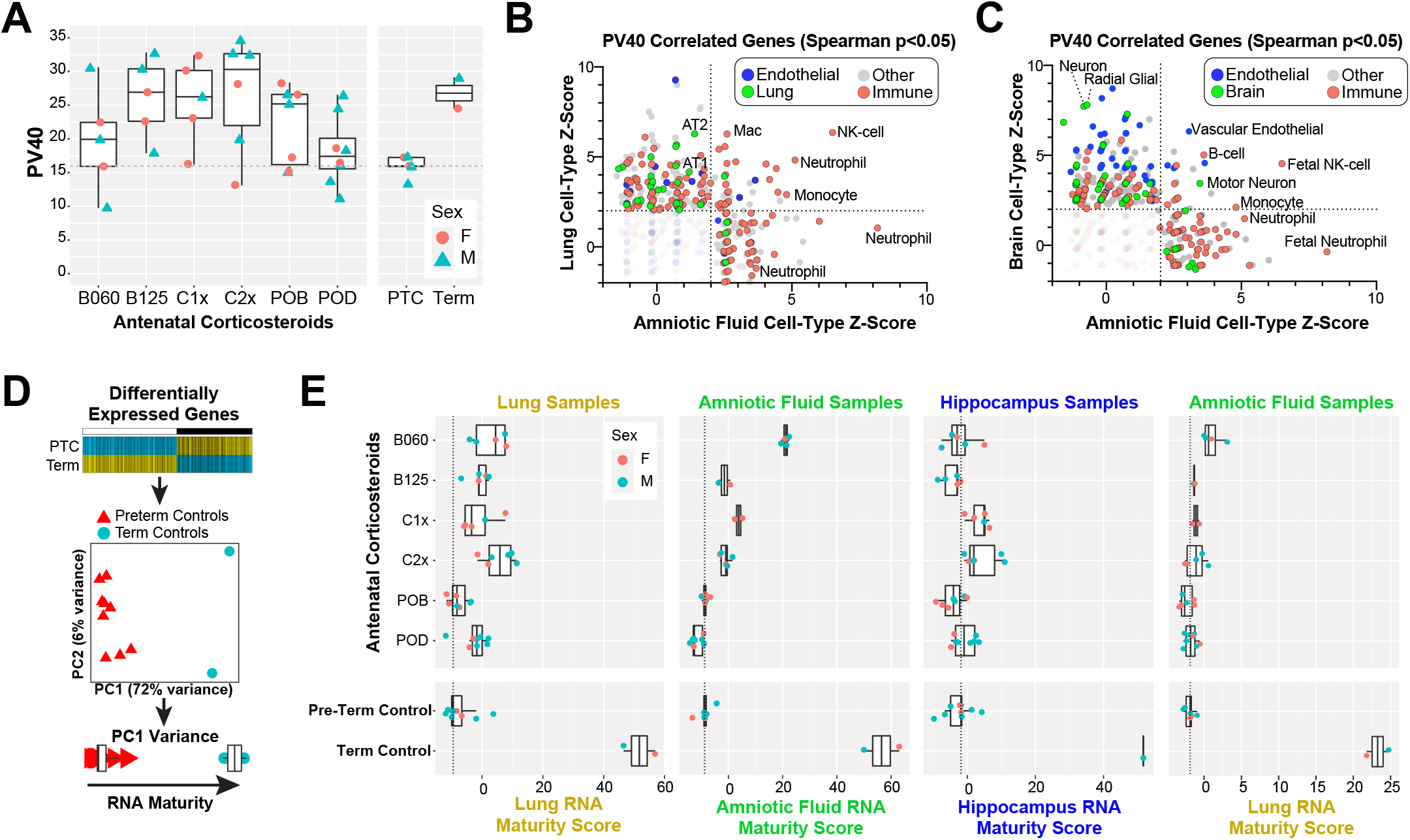
*In Silico* Tissue Maturation Analysis Predicts Optimal Corticoteroid Regimens from Amniotic Fluid. A) Measurement of lung gas volume at 40 cmH_2_O on the pressure volume curve (PV40) was performed on fetal lungs immediately following euthanization. Preterm samples had the lowest P40 and Term samples the highest, with substantial variation among antenatal corticosteroid treated animals. Male and Female fetuses are separately indicated. B) Statistical enrichment of gene-sets from diverse single-cell datasets (n>2,000) for Lung and Amniotic Fluid transcripts that are positively correlated with PV40, across the collection of Rhesus samples (two outlier animals removed). Cell-type signatures associated with Endothelial cells, Lung cells, Immune and other are highlighted, along with significant enrichment results (Z-Score > 2) in either or both Lungs and Amniotic Fluid. C) Same plot as in B, comparing statistically enriched single-cell signatures from brain and Amniotic Fluid samples. D) RNA maturity scoring algorithm design. Differentially expressed genes for Term versus Pre-Term control animals are calculated (see Methods), and selected for PCA of Term and Preterm samples to capture the loadings for PC1 to compute scores for all samples using the PCATools predict function. E) *In silico* maturity scores from each fetus Lung, Brain or Amniotic fluid sample are displayed according to treatment group using either the scoring schema from that same tissue (left three plots) or for Amniotic Fluid scored based on the Lung scoring schema for lung PC1 loading genes.

To determine the prognostic utility of our identified gestation-associated gene signatures, we developed a principal component-based scoring method, leveraging tissue-specific genes that differentiate term and preterm control samples (**Fig. 4D**). Specifically, this method considers RNA-based maturity as a function of the first principal component-associated variance from a PCA analysis. The associated loading genes were then used to project the antenatal corticosteroid treated samples into the established PCA space using the predict function in the software PCATools (**Fig. 4E**). Comparison of lung maturity using the lung RNA Maturity Model (RMM), found that the majority of antenatal corticosteroid treated animals (27 out of 36), had a higher predicted maturity than the upper quartile range of preterm controls, with a Spearman (partial, year) correlation of 0.35 (p=0.023) between the RMM and PV40. Similar to the PV40 data, the lung RMM predicted higher doses of IM Celestone® (1:1 mixture of betamethasone-acetate and betamethasone-phosphate) to have the most significant lung maturation score (Dunnett contrast p-value < 0.001) (**Supplemental Table 10**). Application of the independent amniotic fluid RMM to the treatment cohort, produced results consistent with the lung RMM (Spearman partial (year) rho = 0.45; p=0.008) and found that both lung and amniotic fluid RMM predicted Celestone® and IM betamethasone-acetate treatments to result in the greatest maturation benefit (Dunnett contrast p-value < 0.001), with lower predictions for both oral antenatal corticosteroid regimens (PO betamethasone-phosphate and PO dexamethasone-phosphate). Although lung function should not have an appreciable impact on brain maturity, when the independent Hippocampus RMM was applied to all hippocampus samples, we found Celestone® treatments to result in a moderate improvement in maturity (Dunnett contrast p-value < 0.05), similar to the tissue models. Finally, to determine if the lung RMM was applicable to amniotic fluid samples, we applied the lung model to all amniotic fluid samples, including term and preterm. While it was not inherent that such a model would correctly assign term and preterm samples as the most mature and immature samples, respectively, we did indeed obtain this result (**Fig. 4E right panel, Supplemental Figure 4**). While we observe an improved maturation prediction by IM betamethasone-acetate in the amniotic fluid versus lung, we attribute such this effect to the broader transcriptomic impact of this treatment in the Amniotic fluid, resulting in a greater number of term-induced genes (**Fig. 1D**). Furthermore, while all preterm samples displayed a more compressed dynamic range, more similar to each other, both Celestone® and IM Betamethasone-acetate protocols, were again predicted to provide the greatest benefit, further suggesting that RNAs present in both lung and amniotic fluid can serve as precision markers of lung function and organ maturation.

### Target and side effects of antenatal corticosteroids on amniotic fluid, lung and hippocampus transcriptome

To explore the induction of maturational signaling by antenatal corticosteroids, we identified overlapping genes between term and antenatal corticosteroid therapies when each are compared to preterm controls. While in lung, we frequently observe hundreds of commonly up- and down-regulated genes with different antenatal corticosteroid treatment and term, the total number or percentage of overlapping genes did not correlate with the total corticosteroid dosing (**Supplemental Fig. 5A**). Similarly, in amniotic fluid we find hundreds of common upregulated transcripts in Celestone® term or IM betamethasone-acetate term, these treatments did not correlate with dosing, but do correlate with the RMM predictions (**Supplemental Fig. 5B**). In the brain, we observe far fewer commonly regulated treatment and term transcripts, with relatively few upregulated genes and a greater, but still modest, number of down-regulated genes (less than 106) (**Fig. 5A**).

**Figure 5.**
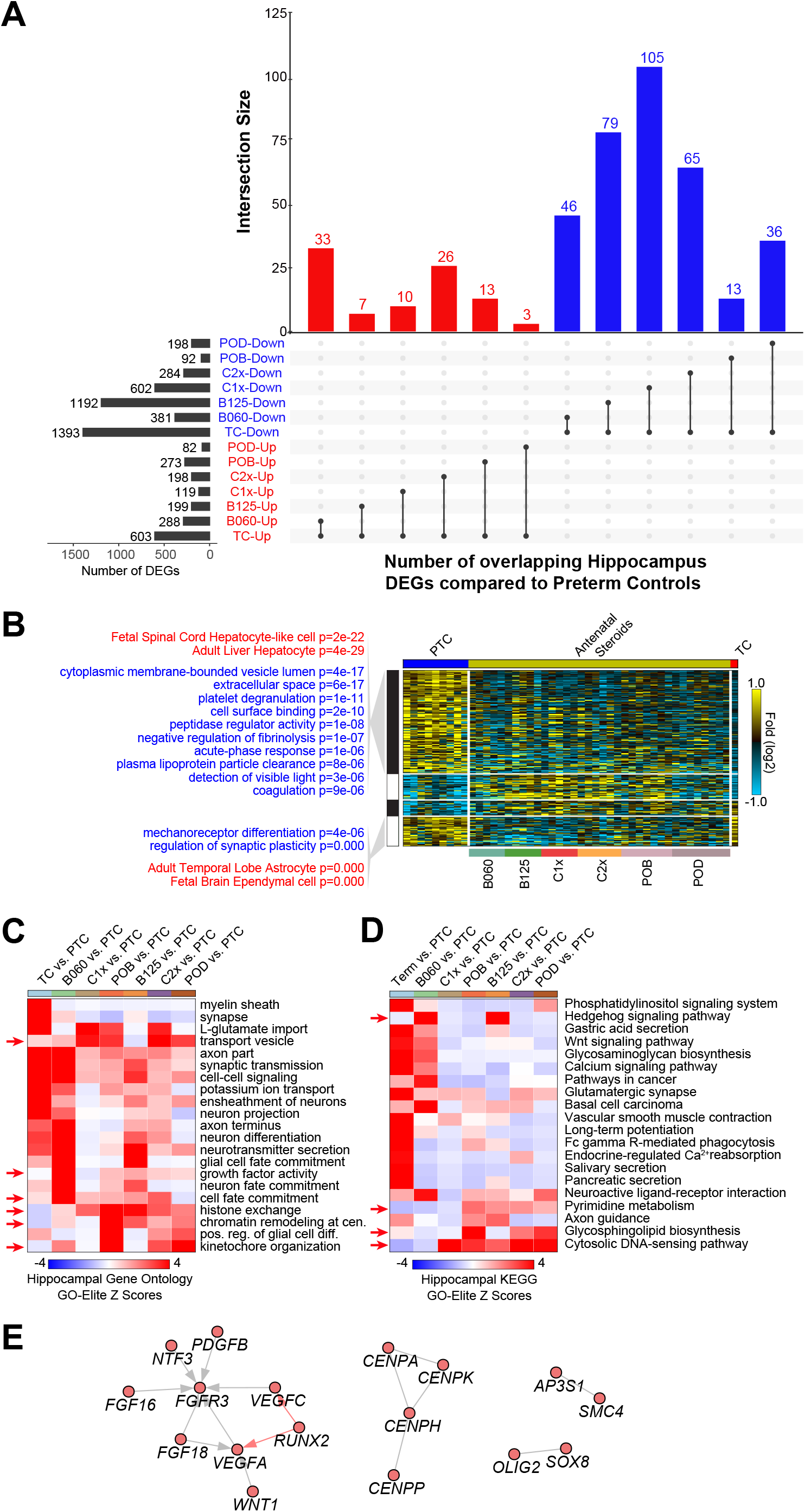
Treatment-specific and common pathways impacted in Rhesus brain. A) Upset plot of overlapping genes in Rhesus term versus preterm hippocampus with genes differentially expressed with each antenatal corticosteroid treatment versus preterm hippocampus. B) Heatmap of predicted marker genes (MarkerFinder algorithm), which are differentially expressed in the union of all antenatal corticosteroid treated samples versus preterm controls. GO-Elite enriched gene sets for Gene Ontology terms (blue) or single-cell RNA-Seq signatures (AltAnalyze BioMarker database) are shown in red, with the corresponding enrichment Fischer’s Exact p-value. C-D) Visualization of the predominant pathways (Gene Ontology (C) and KEGG (D)) observed in term hippocampus or treatment differentially expressed genes versus preterm hippocampus controls (fold ≥ 1.25 and limma p-value ≤ 0.05 adjusted for treatment year effects). The results are displayed as enrichment heatmaps (Z-scores) from GO-Elite. E) Interaction network of upregulated genes associated uniquely differentially expressed in antenatal corticosteroid treatment in the hippocampus. Grey edges represent protein-protein interactions from the BioGRID, WikiPathways or KEGG database, whereas red edges denote predicted transcriptional regulation (PAZAR database). The network was produced using the NetPerspective module of AltAnalyze.

Given the large observed changes in gene expression with corticosteroid therapy (**Fig. 1D**), we asked what were the predominant common and distinct impacted biological pathways and cell-types impacted by antenatal corticosteroids. In lung, we found genes associated with renin-angiotensin system, leukocyte trans-endothelial migration, MAPK signaling pathway, and p53 signaling pathway (Gene Ontology), to be the most frequently shared enriched terms in term and treatment comparisons (**Supplemental Fig. 5C**). When considering scRNA-Seq signatures from human lung, we similarly found shared enrichment in cell types critical for lung maturation and function (AT2, AT1, Neutrophil, dendritic, monocyte, macrophage, NK, mast, basophil) (**Supplemental Fig. 5D**). However, we did not observe frequent enrichment of gene sets restricted to antenatal corticosteroid treated groups. In the hippocampus, we found hundreds of genes differentially expressed in preterm antenatal corticosteroid treated animals versus preterm controls (**Fig. 5C, Supplemental Table 11**). These genes could be separated into the following categories: 1) genes downregulated in treated animals and at term (∼50%), 2) genes downregulated in treated animals but not at term, 3) genes upregulated in treated animals and at term and 4) genes unique upregulated in treated animals. Genes in category 1 were associated with pathways without clear implications for neurodevelopment, whereas genes in category 2 were clearly associated with synaptic transmission, astrocyte identity and mechanoreceptor development, representing likely candidates for deleterious neurological impacts. Focusing on genes upregulated in each of the antenatal treatments but not necessarily shared among the different treatments, we next examined distinct shared pathway impacts. Among enriched KEGG pathway database terms, shared among antenatal corticosteroids but not in term, we observe cytosolic DNA-sensing pathway, glycosphingolipid biosynthesis, pyrimidine metabolism and hedgehog signaling which meet this criterion (**Fig. 5D**). Among enriched Gene Ontology terms, we find histone exchange, chromatin remodeling at centromere, transport vesicle, kinetochore organization and cell fate commitment (**Fig. 5E**). Network analysis of genes associated with these antenatal corticosteroid induced genes revealed that the most interconnected nodes were those involved in FGF and VEGF signaling, oligodendrocyte identity (OLIG2, SOX8) and centromere organization (**Fig. 5F**). Hence, we identified gene networks impinging upon synaptic transmission and glial maturation to be dysregulated with antenatal corticosteroid treatment.

## Discussion

Fetal maturity testing has been limited by the lack of accuracy and markers for global fetal assessment beyond the lung, although it is still used by most obstetricians and maternal-fetal medicine specialists (Visconti et al., 2018). Here we provide proof-of-concept that amniotic fluid contains fetal maturity markers indicating normal gestational advancement as well as distinct changes due to corticosteroid treatment. With advancement of gestational age from 80% to near-term, we found that there are significant changes in the amniotic fluid transcriptome that correlate with transcriptomic changes in the fetal lung and the fetal hippocampus in rhesus macaques. In addition, the transcriptomic effects of antenatal corticosteroid treatment on the lung and hippocampus overlap with differences in the amniotic fluid transcriptome. These findings have important implications for future research and clinical practice. First, we can use screening of the amniotic fluid, which has a larger proportion of cell-free fetal RNA and DNA, to detect organ-specific fetal maturation biomarkers. The eventual goal will be to develop a less invasive test on maternal serum to search for these identified biomarkers. In addition, we can use transcriptomic changes in the amniotic fluid to detect tissue specific effects of antenatal exposures, such as corticosteroids. These findings could lead to new methods for detecting effects of antenatal exposures in humans and guiding the delivery of complicated pregnancies.

We have focused on the fetal lung and hippocampus because of the relevance those organs have for short-and long-term outcomes of preterm infants. Historically, lung immaturity has been a major obstacle to the survival of preterm infants and today, pulmonary complications continue to be the most common morbidity among preterm infants (Jobe and Ikegami, 1998; Stoll et al., 2015). As the survival of preterm infants has improved over the last several decades, focus has shifted to the improvement of neurodevelopmental outcomes of extremely preterm infants (Adams-Chapman et al., 2018; Woythaler, 2019). Despite the established beneficial effects of antenatal corticosteroids in improving the survival of these infants, concerns regarding their safety remain due to the unknown effects on long-term neurodevelopmental outcomes, especially among late preterm pregnancies and the 40% of corticosteroid-treated pregnancies that do not deliver preterm but unnecessarily received the intervention (Jobe and Goldenberg, 2018; Melamed et al., 2019; Paules et al., 2017). During fetal life, the hippocampus is rich in glucocorticoid receptors, and concerns exist about the detrimental effects of antenatal corticosteroids to brain development. Specifically, increased apoptosis in the hippocampus of rhesus macaques was noted (Uno et al., 1994), which is consistent with findings of decreased neuronal density of human newborns and suppressed neuronal development signaling in rhesus macaques exposed to antenatal corticosteroids (Schmidt et al., 2019b; Tijsseling et al., 2012).

Obtaining tissue-specific information on fetal maturation could allow for personalized prenatal and postnatal care of preterm infants. Such individual-specific information could tailor the use of antenatal therapies, such as antenatal corticosteroids or magnesium, to improve neonatal outcomes, and improve the ability to detect patients that would benefit from other prenatal interventions in future trials. Moreover, as our understanding on the dosing of antenatal corticosteroids improves, it could provide additional guidance on the formulation, dosing and length of therapy (Jobe and Goldenberg, 2018; Kemp et al., 2019). It could also help detect specific pathways of beneficial and detrimental effects of antenatal corticosteroids to the fetus and identify surrogate markers for use in clinical trials for new indications of this therapy such as late preterm deliveries or prior to elective cesarean. Based on the results from our in silico predictive model, amniotic fluid generally represents a proxy for the prediction of lung maturation. However, we note greater gene expression impacts for some treatments in the amniotic fluid, compared to the lung, suggesting possible gene expression impacts in non-lung tissue that may contribute to these scores. As such, these molecular differences may not directly relate to maturity, lung compliance or other clinical outcomes that correlate.

One of the limitations of this study is the timing of tissue collection on animals treated with antenatal corticosteroids. There is a disconnect between the timing of peak RNA changes induced by antenatal corticosteroids, which happens early after administration of the drugs, and the timing of measurable fetal lung maturation, which happens after the RNA signaling has decreased (Schmidt et al., 2019a). Hence, it is possible that if measured early after administration of antenatal corticosteroids, the overlap of differentially expressed genes between fetal tissue and the amniotic fluid would be greater than what we reported, and we would be unable to accurately report fetal lung maturation induced by the treatment.

We found that cell-free fetal RNA in the amniotic fluid correlates with tissue specific transcriptomic changes and provides a platform for discovery of fetal maturation and health markers. Our findings need to be further corroborated in human cohorts, not only at different gestational ages but also after fetal treatments, for similarities in gene expression patterns in the amniotic fluid and correlation with clinical outcomes of newborns. Further investigation of cell-free RNA in the maternal serum could provide a less invasive, alternative test for fetal maturation and well-being.

## Methods

### Animals

Pregnant rhesus macaques were studied at the California National Primate Research Center at the University of California Davis with protocols and procedures approved by the Institutional Animal Care and Use Committee. Time-mated pregnant Rhesus macaques were delivered preterm at 132 days of gestation or near term at 155 days of gestation (full term is 165 days). For antenatal corticosteroid dosing studies, preterm pregnant rhesus macaques received one of the treatments listed (Table 1). Oral drugs were given to animals habituated to receiving small treats as previously reported (Schmidt et al., 2019b). The IM betamethasone-acetate 0.125mg/kg and the oral betamethasone-phosphate have been previously shown to induce fetal lung maturation similar to the standard clinical treatment in preterm rhesus macaques, while the betamethasone-acetate 0.06mg/kg and the oral dexamethasone-phosphate did not (Schmidt et al., 2019a; Schmidt et al., 2019b).

Fetuses were delivered 5 days after the initiation of treatment with antenatal corticosteroids at 132 ± 2 days of gestation by cesarean section with intact membranes (n=5-8 animals/group). After delivery, the amniotic fluid was sampled and immediately stored in AssayAssure tubes with standardized buffer (Sierra Molecular Corporation, Incline Village, NV) and frozen at -80 °C. After euthanasia, pressure-volume curves were measured by inflating lungs to 40 cmH_2_O and followed by deflation with measurements of lung volumes using a syringe and pressure manometer. Samples from the right lower lobe of fetal lung and the hippocampus were snap frozen for RNA-sequencing.

### RNA isolation and sequencing

Approximately 10 mL of amniotic fluid from each animal was used for cell-free RNA extraction with the QIAamp Circulating Nucleic Acid Kit (QIAGEN, Valencia, CA). RNA from the lung and hippocampus was extracted with the RNeasy Universal Mini Kit (QIAGEN, Valencia, CA). The RNAseq libraries were prepared using the TruSeq RNA Library Prep Kit v2 (Illumina) with paired-end 75 bp RNA-Seq performed with a HiSeq 2500 (Illumina). Samples were sequenced at a target depth of 30 million reads by the Cincinnati Children’s Hospital Sequencing Core. The sequencing data has been deposited in GEO (GSE171669).

### RNA-Seq analysis quantification

We combined the raw RNA-Seq data from the current study (n=110) with lung and brain samples sequenced in our prior study (n=18, preterm control and antenatal steroids), using the identical experimental protocols (Schmidt et al., 2019a; Schmidt et al., 2019b). To quantify gene expression, we applied pseudoalignment with the software Kallisto using the Ensembl 91 Rhesus Macaque reference transcriptome, using the software AltAnalyze version 2.1.4. For downstream-analyses, Rhesus gene symbols were converted to human gene symbols, where possible (Ensembl BioMart). To correct for seasonal differences in RNA collection and processing, fetal sex and treatment effect, we performed batch effects correction using NOISeq package in R, after excluding genes with strong sex-linked profiles (n=17). NOISeq was run using the ARSyNseq correction function, specifying the treatment group variable rather than a specific set of batch effect factors due to the number and inter-relatedness of candidate factors. Models fit for differential gene expression analyses included blocking by sample year, dichotomized to the earliest time-point group vs. other, to correspond to the two main treatment-by-year subsets. This ensured treatments were compared to (reasonably) contemporary pre-term controls and provided additional control for any year effects not removed in prior steps. Data from one amniotic fluid (AF) sample was not used because the distribution of expression values for this sample was markedly lower than for the other samples. Gene filtering was done on the basis of mean log2 expression values across samples, separately for Lung, Brain and amniotic fluid. Genes were retained for differential expression and correlation analyses if mean log2 expression values (prior to batch correction) were ≥ 1.0 TPM. Principal component analyses (PCA) were conducted using the pca function supplied in the PCAtools R package. The center & scale options were set to “TRUE”.

### RNA-Seq correlation analyses

Correlation of gene expression with PV40 measurements was quantified, by sample type, by Spearman’s partial correlation coefficient, with sample year (dichotomized to 2018 vs. other) as the control variable. Two preterm control samples with implausibly high PV40 values for a preterm fetus were omitted from this analysis. Computations, including calculation of p-values, were done using the pcor.test supplied in the ppcor R package. Correlations of gene expression between Lung and amniotic fluid sample pairs and hippocampus and amniotic fluid sample pairs were computed using the same methods described above. A total of 35 animals had both Lung and AF samples; 34 had both Brain and amniotic fluid samples. After applying the gene filtering described above and merging of the sample data, correlations were calculated for a total of 7,380 genes in the Lung/AF analysis and 7,198 genes in the Brain/AF analysis. Results were displayed in volcano plots with coloring and labelling of points done on the basis of a p-value [unadjusted p <= 0.05] criterion.

### Differential gene expression

Differential gene expression was evaluated using the limma R package applied to the batch-corrected log2(1+TPM) expression values and was done separately for each sample type. The model incorporated blocking of treatments within sample year (dichotomized to 2018 vs. other) to correspond to the two main treatment-by-year subsets. The trend=TRUE and robust=TRUE variance estimation parameters were used in the call to the eBayes function. DEGs from the limma analyses described above, meeting raw p-value <= 0.05 and absolute value of log2 fold change >= log2(1.25) were displayed in barcharts and volcano plots. Upset-type plots were also prepared to show intersections of DEGs. Note that totals for the combination indicated rather than of DEGs unique to each particular combination. Output from the R package UpSetR (upset function) was used to make the bottom of the plot and the R package ggplot2 was used to create the vertical bars with counts on the top portion of the plot. Gene set enrichment analyses were performed in the software GO-Elite, through AltAnalyze (Emig et al., 2010; Zambon et al., 2012). Prior to enrichment, separate gene lists were created for positive and negative log-fold-changes/correlations with filtering criteria of unadjusted p-value <= 0.05 and additionally abs(log2(fold-change)) >= log2(1.25) for results from DEG analyses. The MarkerFinder algorithm was used to provide additional treatment group based DEG analyses and assessment including production of heatmap displays from AltAnalyze.

### RNA Maturity Modeling

A maturation measure was developed for each sample type by applying the following steps: A) PCA reduction was estimated for a reduced dataset limited to preterm control and term control samples and genes that satisfied the following criteria applied to the results of the DEG analyses described above - Lung: abs(log2(fold change)) ≥ log2(2) and fdr-adjusted p-value ≤ 0.05, Brain: abs(log2(fold change)) ≥ log2(1.5) and fdr-adjusted p-value ≤ 0.05, amniotic fluid : abs(log2(fold change)) ≥ log2(1.5) and fdr-adjusted p-value ≤ 0.05, B) Data from all samples were centered and scaled (by gene) and scored on the 1^st^ PC from step A) using the PCATools predict function, C) Sample scores were displayed by treatment in a boxplot graphic. In order to determine the extent to which Lung and Brain maturation might be reflected in the amniotic fluid samples, the Lung and Brain measures were also used to score the AF data. AF sample scores were calculated and displayed following steps B) and C) above using batch-corrected but unfiltered expression values. The lung maturation scale values (as applied to lung data) were plotted (ggplot) versus available PV40 values. A loess smooth function (red dashed line) reflecting the association was fit and displayed (ggplot smooth function). The Spearman (rank) correlation coefficient, adjusted for sample year as described above), was calculated with accompanying p-value using the R package ppcor (pcor.test function). To test for differences in the RNA Maturity Metric between each treatment or term controls versus preterm controls, we performed an Analysis of Variance followed by a Dunnett’s test to obtain a t-statistic and associated p-value for each pairwise comparison. The associated R scripts for the RNA Maturity Metric is available at: https://github.com/DanSchnell/DS_Bioinformatics/tree/Rh_manuscript

### Bulk RNA-Seq Deconvolution Analysis

To estimate the cellular composition of the Lung, Brain and amniotic fluid samples, we applied the DeconRNASeq R package, which uses nonnegative quadratic programming to assess the relative contribution of different cell-types to a bulk sample, against established single-cell populations or bulk-RNA-Seq tissue compendium samples sets. Prior to analysis, we downloaded the already processed sparse-matrix counts for a recently published human lung single-nucleus RNA-Seq dataset spanning neonates through adults (Wang et al., 2020) from LungMAP (GSE161381) and a brain single-nucleus RNA-Seq dataset of pediatric and adolescent neocortex (Velmeshev et al., 2019). Gene UMI counts were scaled to the total reads per nuclei-barcode (counts per 10,000) and collapsed to centroids (pseudo-bulk) profiles for each author annotated nuclei-population. Separately, FASTQ files were downloaded from a prior bulk RNA-Seq study profiling 64-tissue regions in healthy baboons (Mure et al., 2018) (GSE98965). This raw data was aligned to the Papio anubis genome (Panu_3.0) using STAR, with gene expression quantified from exon-exon junction reads in AltAnalyze version 2.1.3 (EnsMart96 database). Gene expression was averaged (centroids) for each tissue for all biological replicate samples. For DeconRNASeq, the top 200 maker genes (MarkerFinder algorithm, AltAnalyze) were obtained for prior to population averaging, for each cell population/tissue to optimal deconvolution. Batch corrected Rhesus TPM values were supplied to DeconRNASeq using the default parameters.

## Acknowledgements

This work was funded by the Cincinnati Children’s Hospital Perinatal Institute Pilot and Feasibility Award (B.D.K., A.F.S.), the National Institute of Child Health and Human Development (HD096256-01A1, B.D.K., A.H.J., N.S., D.T.S.), the National Heart, Lung, and Blood Institute (U24HL148865-02, N.S.), the Bill & Melinda Gates Foundation (OPP1132910, A.H.J.), the National Institute of Environmental Health Services (U01ES029234, C.A.C.), and the Cincinnati Children’s Hospital Academic and Research Committee Grant (C.A.C.).

## Author contributions

Conceptualization, A.F.S., N.S.; B.D.K.; Methodology A.F.S., A.H.J., N.S., B.D.K.; Formal analysis D.S., K.E., K.C., N.S.; Investigation A.F.S., D.S., K.E., K.C., D.T.S., P.S., L.A.M., A.H.J., N.S; Resources L.A.M, A.H.J., N.S., B.D.K.; Data Curation, D.S., N.S.; Writing – Original draft, A.F.S, N.S; Writing – Review & Editing, A.F.S., D.S., D.T.S., A.H.J., N.S., B.D.K.; Supervision, B.D.K. and N.S.

## Supplemental Figures

**Supplemental Figure 1. Antenatal corticosteroids produce distinct transcriptional responses in fetal amniotic fluid and brain**. A-B) Heatmap of the top most specific marker genes for A) hippocampus or B) amniotic fluid RNA-Seq for each treatment. Expression values are calculated as log2 fold-changes relative to the mean of each row. The top GO-Elite Gene Ontology enrichment results are denoted to the left of each cluster along with corresponding Fischer’s Exact test enrichment p-values.

**Supplemental Figure 2. Developmental programs detected in amniotic fluid by differential expression analysis**. A) Volcano plot of differentially genes comparing the limma p-value and fold-change for all genes in term amniotic fluid versus preterm controls. Prior defined human term induced genes from amniotic fluid, are highlighted red (significant) or blue (non-significant) (fold ≥ 1.25 and limma p-value ≤ 0.05 adjusted for treatment year effects). B) Upset plot of overlapping differentially expressed genes comparing either term lungs to preterm lungs, term amniotic fluid to preterm amniotic fluid or term brain to preterm brain, up-or down-regulated.

**Supplemental Figure 3. Shared gestationally induced genes from lung and amniotic fluid impinge upon immune activation and associated transcription factors**. A-B) GO-Elite enrichment networks, indicating each enriched gene-set (central square node) and shared lung-amniotic fluid upregulated genes (red). Blue nodes denote tissue or cellular markers (AltAnalyze TissueMarker database)(A) and yellow nodes denote putative regulatory transcription factors (B). For predicted transcription factor targets, interactions from the PAZAR and Amadeus databases was used (GO-Elite). For tissue-specific genes, the AltAnalyze TissueMarker database was used, with markers defined using the MarkerFinder algorithm applied to diverse tissues and purified cell-types (Kamath-Rayne et al., 2015).

**Supplemental Figure 4. Correlation of RNA Maturity Metric scores in lung and amniotic fluid**. Scatter plot of computed lung maturity scores using Rhesus lung RNA-Seq with the lung model (x-axis) and amniotic fluid maturity scores from Rhesus amniotic fluid RNA-Seq with the amniotic fluid model (y-axis).

**Supplemental Figure 5. Treatment-specific and common pathways impacted in Rhesus lung and amniotic fluid**. A-B) Upset plot of overlapping differentially expressed genes in different antenatal corticosteroid treatments with Term controls in the A) Rhesus Lungs and B) Rhesus Amniotic Fluid. C-F) Visualization of the predominant pathways (KEGG (C) and Gene Ontology (E)) observed in term or treatment differentially expressed genes versus preterm controls (fold ≥ 1.25 and limma p-value ≤ 0.05 adjusted for treatment year effects). The results are displayed as enrichment heatmaps (Z-scores) from GO-Elite for (C-D) Lung and (E-F) Amniotic Fluid.

