## Supplementary figures and images for "Fetal maturation revealed by amniotic fluid cell-free transcriptome in rhesus macaques"

### Supplemental Figure 1

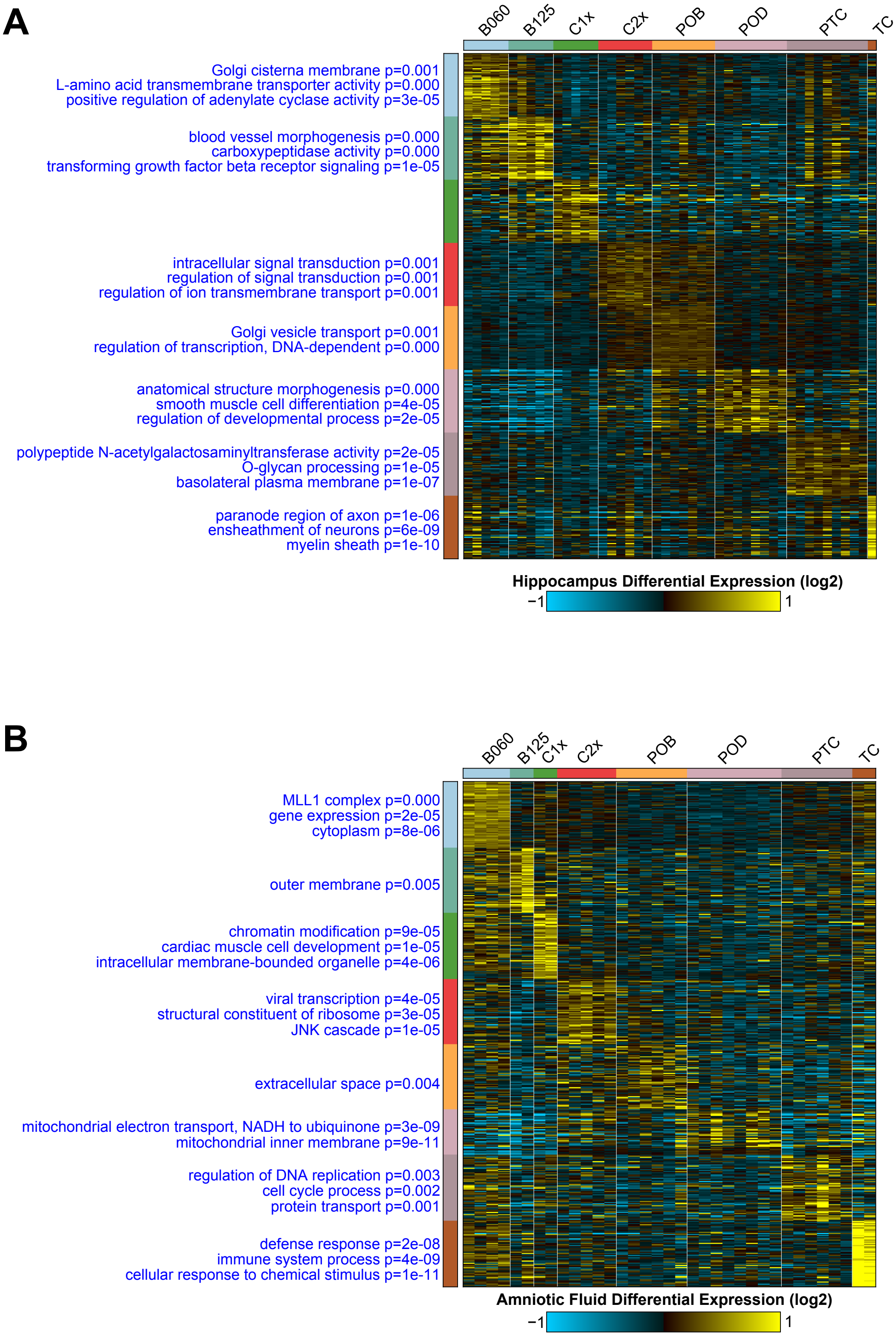


Supplemental Figure 1

### Supplemental Figure 2

A

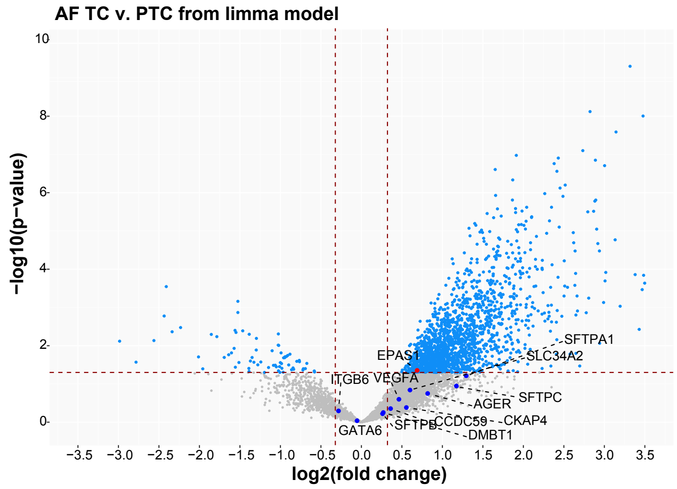


B

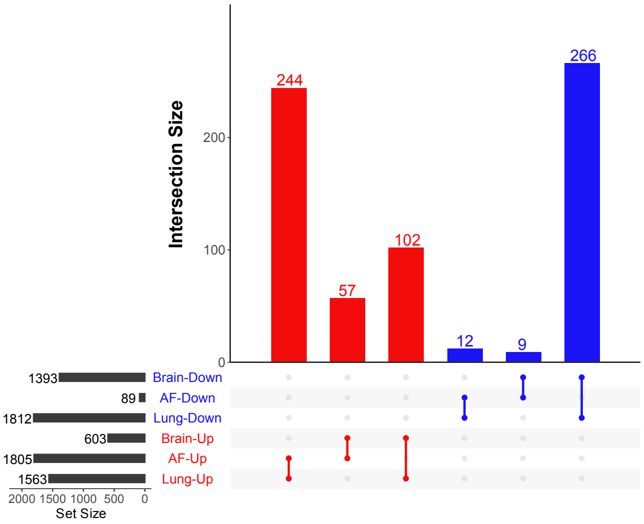


Supplemental Figure 2

### Supplemental Figure 3

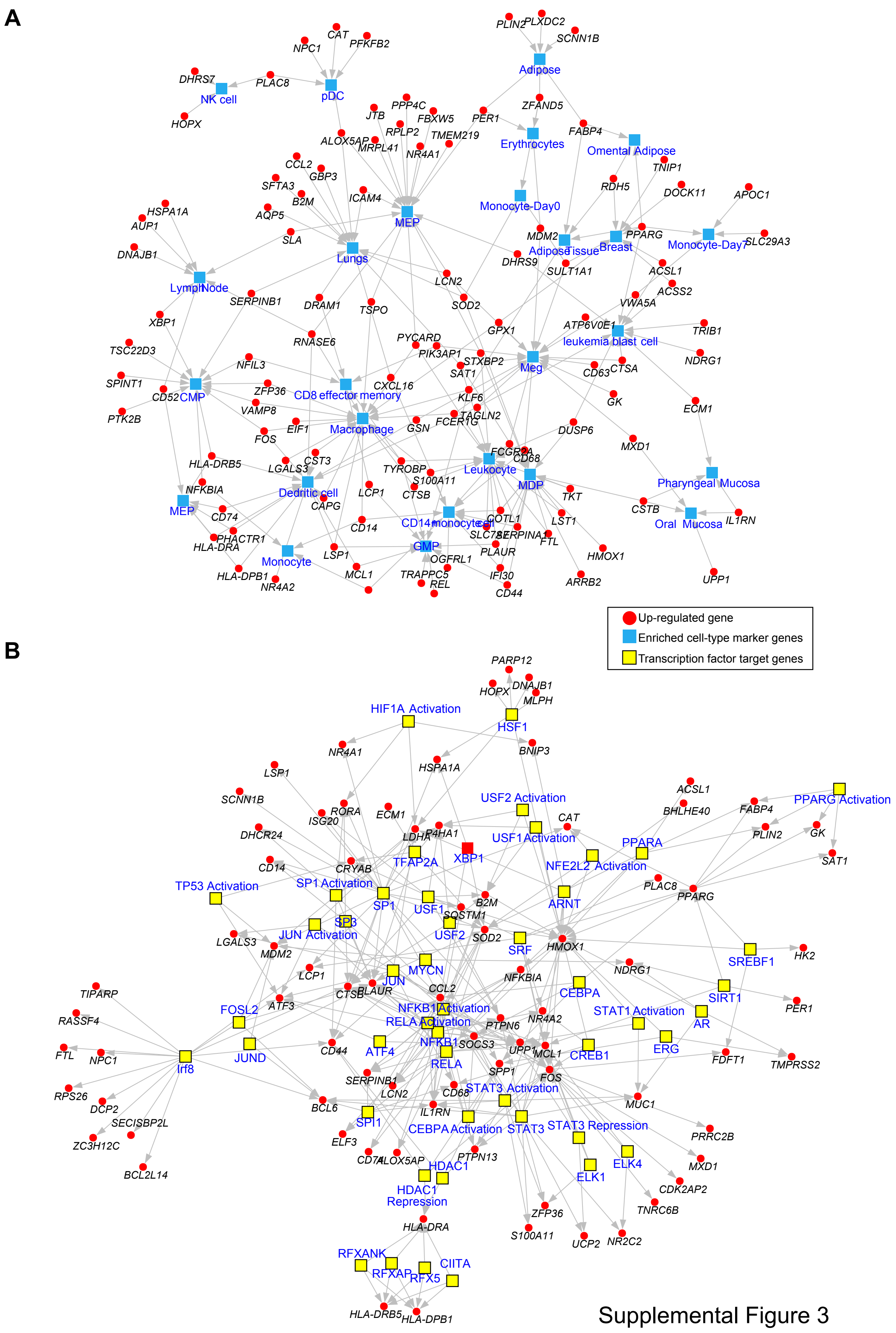


Supplemental Figure 3

### Supplemental Figure 4

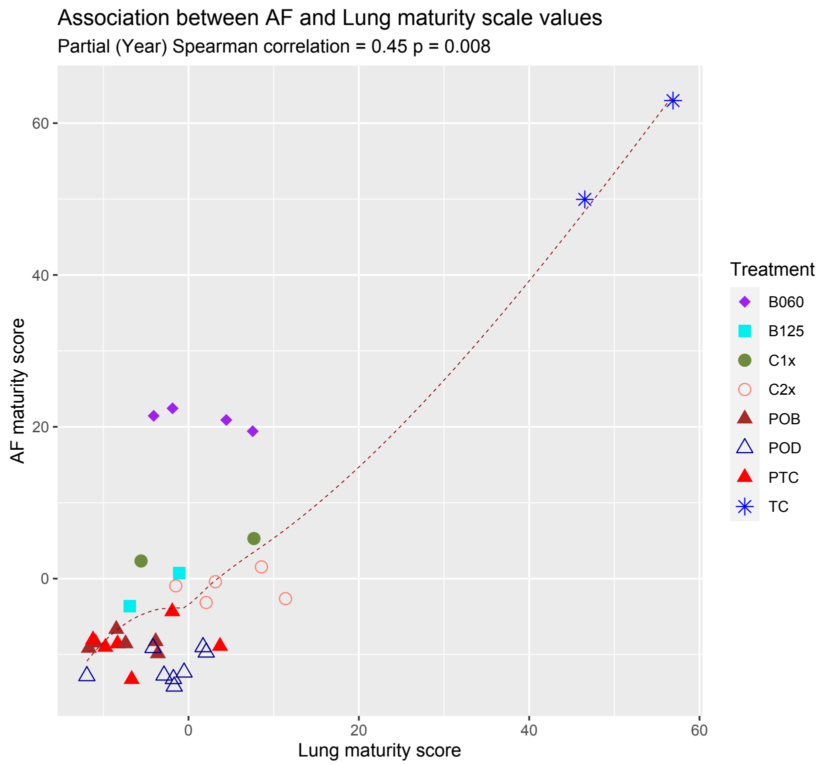


Supplemental Figure 4

### Supplemental Figure 5

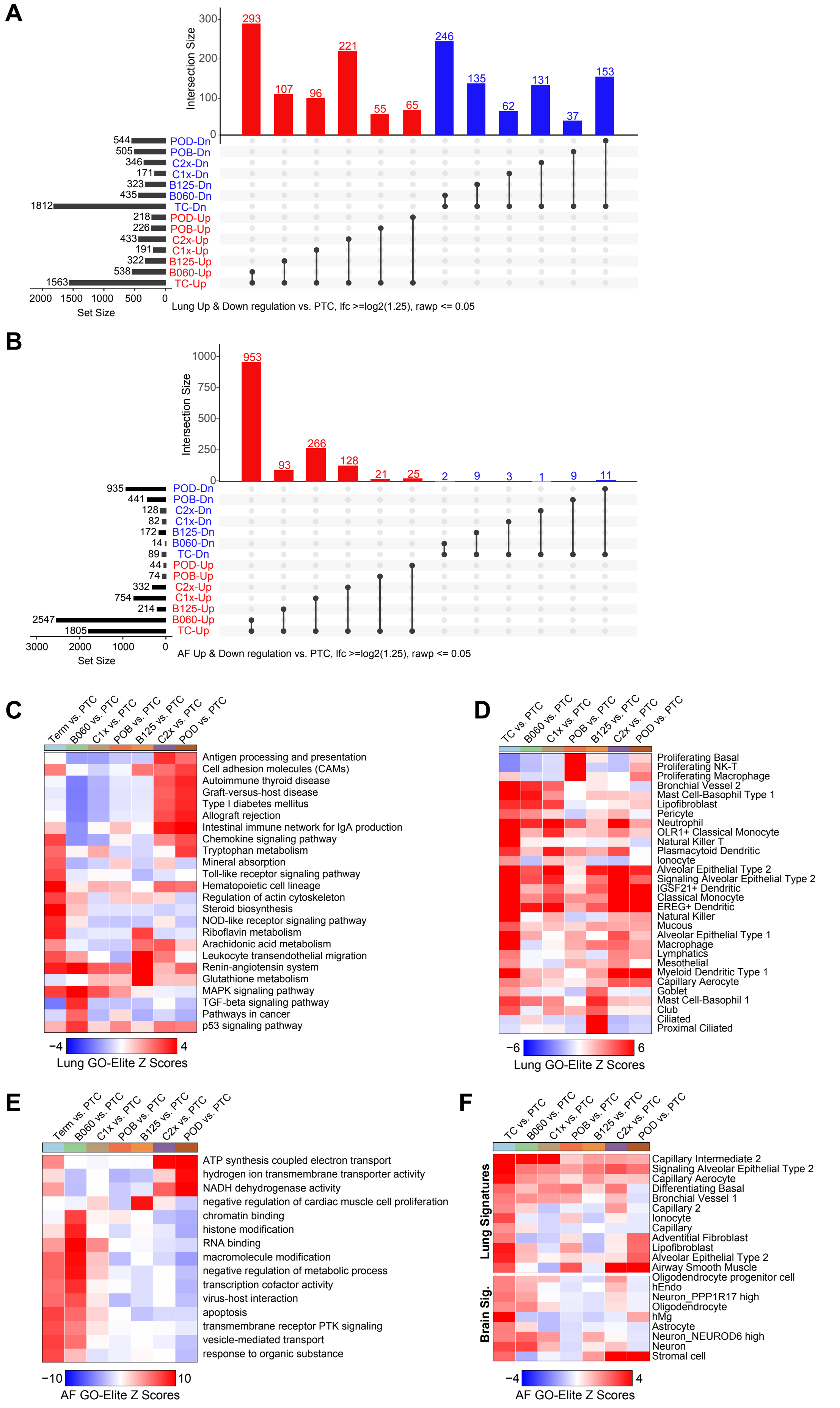


Supplemental Figure 5
